# SOX8-USP7-PGC-1α axis enhances thermogenesis in brown adipocytes

**DOI:** 10.64898/2026.05.22.727126

**Authors:** Yuqin Gu, Ziheng Kan, Guangyu Lu, Yongcong Cai, Xingjing Yang, Qi Zhu, Yuwei Li, Xing He, Xue Yang, Zhenglin Yang, Hao Qian, Ziyan Wang

## Abstract

Obesity is one of the most prevalent diseases worldwide. Increasing thermogenesis to enhance energy expenditure has emerged as a promising therapeutic strategy. In an effort to identify new regulatory targets in thermogenic adipocytes, we found that SOX8 is correlated with obesity and serves as a novel marker of classical brown adipocytes in both humans and mice, upregulating during acute cold exposure. Functional studies further demonstrated that adipocyte-specific knockdown of SOX8 leads to obesity and metabolic dysfunction in mice. Mechanistically, SOX8 directly interacts with USP7 and stabilizes PGC-1α by reducing its K48-linked polyubiquitination. AAV-Rec2-mediated SOX8 overexpression initially enhanced energy expenditure, improved insulin sensitivity, and alleviated metabolic dysfunction in HFD-fed mice. However, prolonged SOX8 overexpression induced compensatory metabolic maladaptation, characterized by reduced energy expenditure, impaired glucose homeostasis, and mitochondrial structural disruption. These findings reveal a novel SOX8–USP7–PGC-1α regulatory axis in brown adipocytes, and reveal a previously unrecognized time-dependent effect of sustained thermogenic activation, highlighting SOX8 as a promising therapeutic target for obesity and metabolic syndrome.

## Introduction

Obesity represents a major global health challenge, characterized by excessive adiposity and associated metabolic dysfunctions, including insulin resistance, dyslipidemia, and systemic inflammation(Blüher, 2019). The imbalance between energy intake and expenditure underlies the accumulation of adipose tissue, promoting the development of metabolic syndrome and increasing the risk of type 2 diabetes, cardiovascular disease, and other obesity-related complications(Perdomo *et al*, 2023). Despite widespread lifestyle and pharmacological interventions, sustainable strategies to prevent or reverse obesity remain limited, highlighting the need for novel approaches targeting energy balance and adipose tissue function(Rosenbaum & Foster, 2023). Understanding the intrinsic regulatory mechanisms that govern adipocyte metabolism is therefore critical to identifying potential therapeutic targets capable of restoring systemic metabolic homeostasis.

Brown and beige adipocytes offer a promising avenue for combating obesity due to their intrinsic capacity for adaptive thermogenesis(Harms & Seale, 2013). Unlike energy-storing white adipocytes, brown adipocytes dissipate chemical energy as heat through uncoupling protein 1 (UCP-1)-mediated mitochondrial proton leak, contributing to basal and inducible energy expenditure(Cohen & Kajimura, 2021). Activation or expansion of brown and beige adipose depots has been shown to enhance glucose clearance, improve lipid mobilization, and increase systemic metabolic flexibility, independent of changes in diet or physical activity(Kajimura *et al*, 2015). These properties position brown fat–targeted therapies as attractive interventions for obesity and metabolic disease. However, effective manipulation of brown adipocytes requires a detailed understanding of the molecular networks that coordinate thermogenic gene expression, mitochondrial integrity, and adipocyte identity(Aquilano *et al*, 2023).

PGC-1α is a master regulator of brown adipocyte thermogenesis and mitochondrial biogenesis(Lin *et al*, 2005b). It functions as a transcriptional coactivator with PRDM16(Seale *et al*, 2007) and PPARγ(Puigserver *et al*, 1998) to upregulate UCP-1. In addition to its role in thermogenic gene activation, PGC-1α coordinates a broad transcriptional network by interacting with transcription factors such as NRF-1, NRF-2, PPARα, ERRα, and Sp1 to drive the expression of nuclear genes involved in respiratory chain function, mitochondrial transcription, translation, DNA replication, protein import, and the assembly of mitochondrial complexes(Scarpulla, 2008). PGC-1α deficiency in whole-body knockout models leads to impaired thermogenesis and whitening of BAT(Lin *et al*, 2004). Brown adipocytes lacking PGC-1α reduced the induction of thermogenic genes(Uldry *et al*, 2006), whereas overexpression in human white adipocytes induces browning(Tiraby *et al*, 2003). These observations highlight PGC-1α as a critical node in adaptive thermogenesis and a potential therapeutic target for obesity intervention.

The activity and stability of PGC-1α are dynamically regulated by multiple post-translational modifications, including phosphorylation, acetylation, SUMOylation, and ubiquitination, which integrate diverse metabolic and environmental signals to fine-tune its protein levels, transcriptional activity, and interactions with other factors(Fernandez-Marcos & Auwerx, 2011). Several E3 ubiquitin ligases, such as RNF2(Sen *et al*, 2011), RNF34(Wei *et al*, 2012) and FBXW7(Trausch-Azar *et al*, 2015), target PGC-1α for proteasomal degradation, thereby limiting its stability and suppressing thermogenic gene programs. Conversely, deubiquitinases such as A20 can remove ubiquitin chains from PGC-1α, protecting it from degradation and maintaining its functional capacity(Bombassaro *et al*, 2018). These reversible modifications allow brown adipocytes to rapidly adapt their thermogenic machinery in a context-dependent manner. Therefore, modulating PGC-1α ubiquitination represents a potential strategy to enhance thermogenesis and combat obesity.

In this study, we identify SOX8 as a key regulator of PGC-1α. In previous studies, global knockout of SOX8 in mice resulted in reduced white adipose tissue (WAT) mass, suggesting its involvement in adipose energy metabolism(Guth *et al*, 2009). We found that SOX8 is a specific marker of brown adipose tissue (BAT) and shows correlation with thermogenesis-related obesity. We established that SOX8 enhances thermogenic gene expression and promotes mitochondrial integrity through simultaneously recruiting the deubiquitinase USP7 to stabilize PGC-1α, preventing its K48-polyubiquitination-mediated degradation. Our findings reveal a novel SOX8–USP7–PGC-1α regulatory axis, providing mechanistic insight into how transcription factors can integrate ubiquitin-mediated proteostasis with metabolic regulation. These results suggest that targeting SOX8 represents a promising strategy to harness brown adipocyte thermogenesis and mitochondrial homeostasis for the treatment of obesity and metabolic disorders.

## Methods

### Clinical sample acquisition of human adipose tissue

Human adipose tissues (obtained from the brown and white adipose tissue of the neck region during thyroidectomy procedures) in this study were obtained from Sichuan Cancer Hospital. All subjects have signed written informed consent forms, and the research protocol has been approved by the UESTC and hospital’s Ethics Committee.

### Mice experiments

All animal experiments were approved by the Ethics Committee of Sichuan Provincial People’s Hospital and University of Electronic Science and Technology of China. SOX8^flox/flox^ and Adipoq-Cre mice were obtained from Cyagen Biosciences Inc. SOX8-AKO mice were generated by crossing SOX8^flox/flox^ mice with Adipoq-Cre transgenic mice, and SOX8^flox/flox^ littermates were used as wild type (WT) controls. Other C57BL/6J mice were purchased from GemPharmatech Co., Ltd. All mice were housed in specific pathogen-free (SPF) facilities under standard humidity and temperature conditions, with a 12-hour light/dark cycle starting at 7:00 AM. For the diet-induced obesity (DIO) study, 8-week-old male WT and AKO mice were fed a high-fat diet (HFD) containing 60% kcal from fat (Research Diets D12492) for 9 weeks.

### Cell culture

HEK 293T cells, originally obtained from ATCC, were purchased from Lookbio (Shanghai, China).

Stromal vascular fractions (SVFs) were isolated from the iBAT of 6-8-week-old male mice. Tissues were digested in 2 mg/mL collagenase type II (Solarbio, C8150) for 30 minutes at 37°C. Cells were filtered through a 50 μm cell strainer, centrifuged at 2000 rpm for 10 minutes, and put into 6-well plates. The medium was refreshed after 48 hours.

Human SVFs from neck brown and white adipose tissue were obtained from biopsy samples collected during thyroidectomy. Tissues were cut into small pieces, placed in 6-well plates and covered under Lysine-coated slides (Solarbio, YA0170) to prevent floating. Human SVFs were cultured in DMEM/F12 containing 10% fetal bovine serum (FBS) and 1% penicillin-streptomycin (PS). Others were cultured in DMEM containing 10% FBS and 1% PS. Cells were immortalized by pLV-EF1A-SV40LT or pLV-EF1A-TERT.

### Plasmids and Lentivirus Packaging

SFB-USP7 plasmid was kindly provided by Peijing Zhang’s lab(Chen *et al*, 2024). Other plasmids were constructed by Youbio Biological Company (Changsha, China). Lentivirus was packaged with PEI, psPAX2, pMD2.G in HEK 293T cells. After 72 hours, virus was collected, centrifuged at 100,000×g for 2 hours and resuspended in 200 μL PBS.

### Packaging, purification and infection of adeno-associated virus (AAV)

The AAV-Rec2 system was utilized for overexpression of SOX8 in adipose tissue. The adeno-associated viruses (AAVs) were packaged and purified by Shandong Weizhen Biological Technology Co., Ltd., including the construction of a mouse Sox8 gene recombinant AAV (rec2) and a control AAV-mCherry. After a 6-hour fasting period, AAVs carrying SOX8 (rAAV-SOX8) were administered via oral gavage to 8-week-old male mice at a dose of 2 × 10^10^ vg in 200 µL PBS per mouse. Littermates receiving an equivalent dose of the empty vector (AAV-mCherry) served as the control group. Administration was repeated every two weeks for a total duration of 6 weeks or 12 weeks.

### Adipogenic Differentiation and Oil red O staining

After reaching confluence, cells were induced with medium containing IBMX, T3, insulin, rosiglitazone, dexamethasone and indomethacin for 2 days, then maintained in medium containing T3, insulin and rosiglitazone for 7 days. Oil red O staining (Servicebio, G1015-100ML) followed the manufacturer’s protocols.

### Biochemical index detection

Serum samples were collected according to experimental requirements and sent to Chengdu Lilai Laboratory for the measurement of total cholesterol (TC), triglycerides (TG), free fatty acids (FFA), high-density lipoprotein (HDL), low-density lipoprotein (LDL-C), and alanine aminotransferase (ALT) levels in mice.

### Histology analysis

Tissue samples of mouse iBAT, epididymal WAT (eWAT), inguinal WAT (iWAT), and livers were fixed in 4% paraformaldehyde and subsequently sent to Servicebio (Wuhan, China) for Hematoxylin and Eosin (H&E) staining and Oil Red O staining. The stained slides were photographed by microscope for further analysis by Image-Pro plus software.

### Glucose and insulin tolerance tests

For the glucose tolerance test (GTT), mice were fasted for 8 hours with access to water only, and received an intraperitoneal injection of D-glucose (2 mg/g body weight; Solarbio, G8150). For the insulin tolerance test (ITT), mice were fasted for 6 hours with access to water and intraperitoneally injected with human insulin (0.125 mg/g body weight; Solarbio, I8830). Blood samples were collected from the tail vein at 0, 15, 30, 45, 60 and 90 minutes post-injection, and blood glucose levels were measured using a glucometer.

### Cold Tolerance Test

On the day of the experiment, mice were fasted (with free access to water) for 6 hours and then transferred to individual cages at 4°C for acute cold exposure for 6 hours. Rectal temperature was measured at designated time points using a rectal thermocouple probe.

### Metabolic cage study

For metabolic studies, mice were individually housed in metabolic cages (Oxymax/CLAMS Metabolic Analysis System) for 24 h with free access to food and water. Oxygen consumption (VO₂), CO₂ production rate (VCO₂), food/water intake, and heat were monitored for 48 h. The ambient temperature was maintained at 22°C, with a 12 h light/12 h dark cycle.

### Transmission electron microscopy (TEM)

Dissected BATs were cut into small pieces and fixed in pre-cooled fixation buffer (2.5% glutaraldehyde in 0.1 M phosphate buffer, pH 7.4), and were subsequently sent to Chengdu Lilai Biotechnology Co., Ltd for TEM according to the standard protocols. Mitochondrial length, quantity, mitochondrial damage, as well as lipid droplet size and number were analyzed based on the TEM images by Image-Pro plus software.

### Western Blotting (WB)

Cells or tissues were lysed in RIPA buffer (Solarbio, R0010) containing protease (Solarbio, P0100) and phosphatase inhibitor cocktail (Solarbio, P1260). The protein concentration was analyzed by BCA Protein Quantitative Kit, and the samples were separated by SDS-PAGE, then immunoblotted with the indicated antibodies. Primary antibodies were diluted as follows: anti-Tag and anti-β-TUBULIN antibodies were used at 1:5000, whereas all other primary antibodies were used at 1:1000.

### RNA extraction and RT-qPCR

Total RNA was extracted with TRIzol (Solarbio, R1100). The cDNA synthesis was conducted using HiScript III qRTSuperMix (Vazyme, R333-01). The qRT-PCR assay was performed with SYBR Green (CWBIO, CW3360M) following the manufacturer’s protocols. A list of the primers (Tsingke, Chengdu, China) used in this study is provided in Supplementary Table S1.

### Mitochondrial Respiration Assay

Mitochondrial respiratory function in cells was assessed using an Oroboros Oxygraph-2k (Innsbruck, Austria) in a temperature-controlled chamber. The cells were counted and seeded into chambers. After equilibration, oligomycin (5 nM), FCCP (0.5 µM), and Antimycin A (2.5 µM) were added sequentially. These inhibitors were obtained from Huawei Corporation (Cat:023004). Data were collected using DatLab acquisition software 5.2 (Oroboros Instruments, Innsbruck, Austria).

### Cycloheximide (CHX) chase assay

SVF-induced mature brown adipocytes were treated with fresh maintenance medium containing 200 μg/mL cycloheximide (CHX), with the time of addition defined as 0 h. Cells were harvested at 0, 2, 4, 8, and 12 h, lysed, and subjected to Western blot analysis to assess PGC-1α or USP7 protein levels.

### Immunofluorescence (IF)

Cells were seeded on coverslips (Biosharp, BS-14-RC). After adipogenic differentiation, cells were fixed with 4% paraformaldehyde for 15 min and permeabilized with 0.1% Triton X-100 for 10 min. After washing three times with PBS, cells were blocked with 5% BSA in PBS for 1 h. Primary antibodies were diluted in 1% BSA and applied to the cells, followed by incubation overnight at 4°C. After three washes with PBS, cells were incubated with secondary antibodies, BODIPY and DAPI for 1h. Finally, coverslips were mounted with anti-fade mounting medium and imaged using confocal microscopy (Zeiss).

### Protein purification

After indicated plasmids transfection by lipo2000 (Thermo, 11668019) in HEK293T, protein was purified with indicated beads at 4℃ for 4 h and eluted with 100 μM S peptides (Beyotime, P9816) or 2mg/mL D-biotin at 4℃ for 30 min.

### Protein-protein binding assay

Antibody-linked magnetic beads were produced as following. Every 2.5 μg antibodies were linked with 25 μL protein A/G magnetic beads (MCE, HY-K0202) by 5 mM BS3 (MCE, HY-124329) for 1 hour at room temperature and stopped by 1 M Tris-HCl, pH=7.5 (Solarbio, T1170) for 15 min.

For endogenous Co-IP, mouse was exposed at 4°C for 6 hours and iBATs were lysed in native lysis buffer (Solarbio, R0030) containing protease and phosphatase inhibitor cocktail. After the protein concentration was measured, every 2 mg protein was pre-cleared by 25 μL protein A/G magnetic beads for 1 hour and immunopreciped by 25 μL antibody-linked magnetic beads at 4°C overnight. After 4-time washing by native lysis buffer, protein was eluted by 25 μL loading buffer (Solarbio, P1040) at 95°C for 10 min, separated by SDS-PAGE and immunoblotted with indicated antibodies.

For exogenous binding assay, plasmids were transfected by lipo2000 in HEK293T. After 12-24h, protein was immunoprecipitated with indicated beads and immunoblotted with indicated antibodies.

For in vitro binding assay, control S-HA peptides and USP7 protein were purified by S-protein agarose from HEK293T transfection lysis at 4°C for 4 h. After 4-time washing, SOX8 purified protein was added and reacted at 4°C overnight. After 4-time washing, protein was eluted by loading buffer, separated by SDS-PAGE and immunoblotted with indicated antibodies.

### Ubiquitination assay

After 6 h 40 μM MG132 treated, iBAs were lysed and immunoprecipitated with anti-PGC-1α or anti-USP7 magnetic beads and immunoblotted with K48 and Ub antibodies.

### In vitro deubiquination assay

After 6 h 40 μM MG132 or CQ treated, protein was purified by indicated beads from HEK293T transfection lysis at 4°C for 4 h. Purified protein was added and reacted with 4 mM DTT at 37°C for 30 min and stopped by loading buffer, separated by SDS-PAGE and immunoblotted with K48 and K63 antibodies.

### Chromatin immunoprecipitation (ChIP)

After adipogenic differentiation, SOX8-OE-iBAs were chromatin-immunoprecipitated by anti-FLAG and control rabbit IgG with ChIP Assay Kit (Beyotime, P2078). DNA was purified with DNA Purification Kit

(Beyotime, D0033). Primers (F1-CCACCTTACAATGTCTGAACTGG, R1-GGACAGAGGGTTGGTTGGCT) (F2-GGGGGTTTATTCTAGCCTCGG, R2-GAAGCTGCAAGACCTATGGATGA) were used for qPCR with SYBR Green.

### Luciferase reporter assay

Immortalized SVFs were transfected with Lipo 8000 (Beyotime, C0533), control Renilla plasmid and pGL3-*UCP-1*-promoter. After 48 h, cells were performed with Dual Luciferase Reporter Gene Assay Kit (Yeasen, 11402ES60). Data were collected using LumiPro (Alpha, Suzhou, China).

### RNA sequencing and bioinformatics analysis

Interscapular brown adipose tissue samples (∼50 mg per sample) were collected and subjected to RNA sequencing by Biomarker Technologies (Beijing, China). RNA extraction, library construction, and sequencing on the Illumina HiSeq platform were performed according to standard protocols. Differentially expressed genes (DEGs) were identified using a threshold of |log₂(fold change)| ≥ 1 and *p* < 0.05.

Publicly available datasets were downloaded from the GEO and GTEX database. Raw data were processed and analyzed using RStudio and relevant bioinformatics packages. After normalization, genes were screened for subsequent analysis.

### Protein–protein interaction structure prediction

Protein sequences of human SOX8 (UniProt ID: P57073), USP7 (UniProt ID: Q93009), and PGC-1α (UniProt ID: Q9UBK2) were obtained from the UniProt database. Protein–protein complex structures were predicted using the AlphaFold3 server. The confidence of predicted interactions was evaluated using the ipTM score (range: 0–1), where >0.5 indicates high confidence and 0.2–0.5 indicates medium confidence. The top-ranked models were downloaded and visualized using PyMOL for structural representation.

### Quantification and statistical analysis

All experiments were performed with a minimum of three replicates or triplicate samples. Data are expressed as means ± Standard Error of the Mean (SEM.). Statistical analyses were conducted using GraphPad Prism software (version 10; GraphPad Software Inc., La Jolla, CA). Comparisons between two groups were evaluated using a two-tailed unpaired Student’s t-test, while differences among multiple groups were assessed via one-way analysis of variance (ANOVA) with Tukey’s or Dunnett’s post multiple comparison test. Pearson correlation coefficient was calculated to address association of two parameters. P-values for all analyses are presented within the respective Fig. panels. A threshold of P < 0.05 was adopted to denote statistical significance.

## Results

### SOX8 is a specific marker of BAT and correlated with thermogenesis-related obesity

In our previous study, SOX8 regulates de novo fatty acid synthesis and ferroptosis in hepatocellular carcinoma(Yang *et al*, 2024a), indicating SOX8’s role in lipid metabolism and mitochondrial quality control. SOX8 is also recognized as a marker of muscle satellite cells and is downregulated during myogenesis(Schmidt *et al*, 2003). Based on BAT and skeletal muscle share a common MYF5⁺ progenitor(Inagaki *et al*, 2016), this suggests that SOX8 expression in BAT may be upregulated to facilitate lineage divergence from muscle. By extending these findings, we aimed to further delineate the SOX8 regulatory network underlying lipid and mitochondrial homeostasis to explore its potential as a therapeutic target for obesity and related metabolic disorders.

To investigate the relationship between SOX8 and obesity, we analyzed *Sox8* expression in iBAT from mice fed a high-fat diet (HFD). *Sox8* mRNA levels were decreased in interscapular brown adipose tissue (iBAT) with increasing in 2 and 4 weeks of HFD feeding (Fig. 1A), suggesting that SOX8 expression is suppressed in the context of diet-induced obesity. Consistently, SOX8, PGC-1α, and UCP-1 protein levels were all significantly reduced in iBAT from obese mice compared with normal diet (ND) controls (Fig. 1B), and *SOX8* mRNA levels were positively correlated with *PGC-1α* expression in human adipose tissue (Fig. 1C), suggesting that SOX8 may regulate thermogenesis through the PGC-1α pathway.

**Fig. 1.**
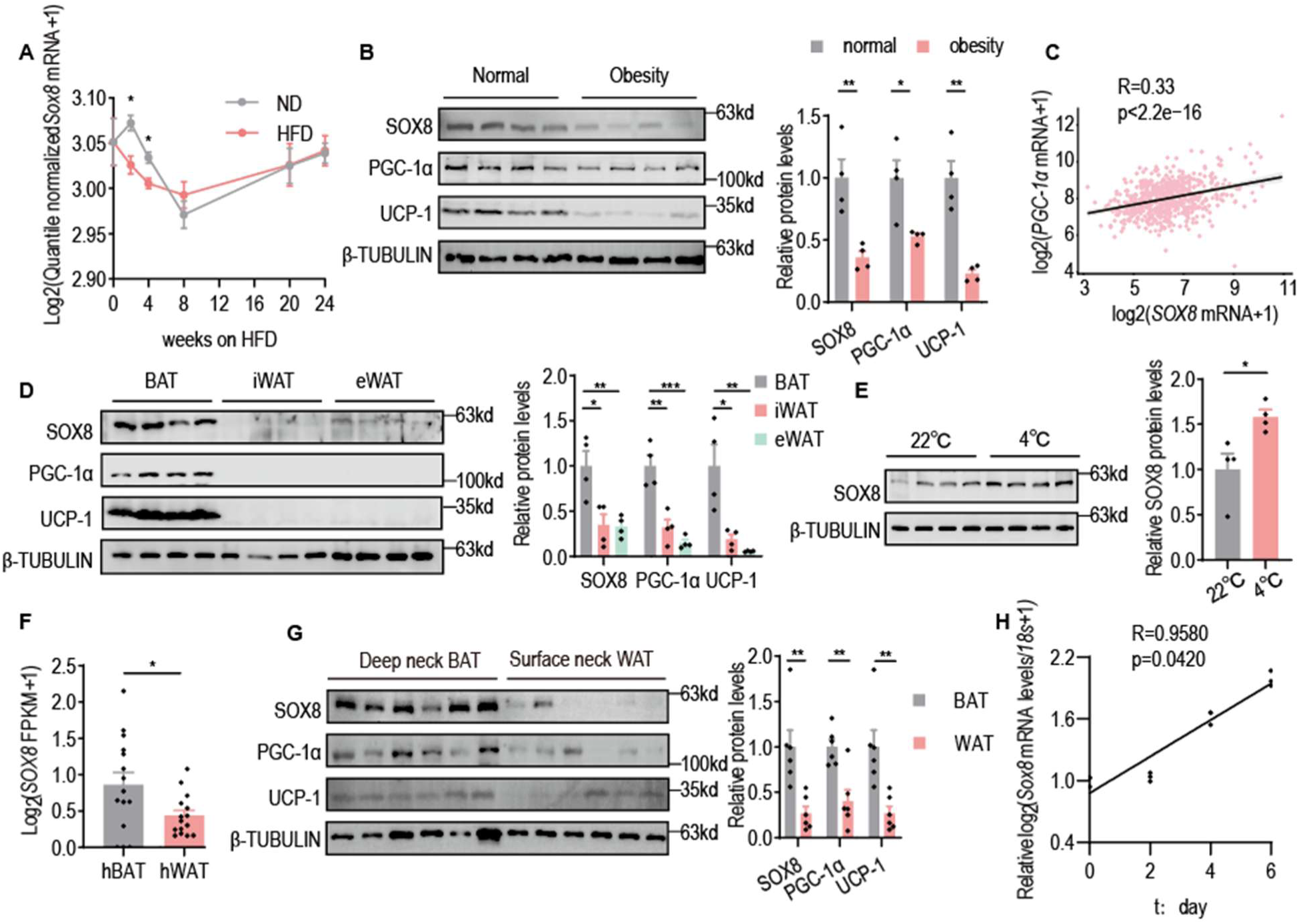
SOX8 is a specific marker of BAT and correlated with thermogenesis-related obesity.[](A) Sox8 mRNA levels in iBAT from ND- and HFD-fed mice at indicated time points. Data were obtained from GSE64718.[](B) Representative blots and quantification of SOX8, PGC-1α, and UCP-1 protein in mouse iBAT from ND and HFD groups (n = 4).[](C) Correlation analysis between SOX8 and PGC-1α mRNA levels in human adipose tissue. Data were obtained from GTEx: SRP012682.[](D) Representative blots and quantification of SOX8, PGC-1α, and UCP-1 protein levels in iBAT, iWAT, and eWAT from mice (n = 4).[](E) Representative blots and quantification of SOX8 protein in mouse iBAT after cold exposure. Mice were fasted for 6 h and then exposed to 4°C for 6 h (n = 4).[](F) SOX8 mRNA levels in human brown adipose tissue (hBAT) and white adipose tissue (hWAT). Samples were re-clustered based on global transcriptomic profiles using Z-score normalization and high thermogenic gene expression were defined as BAT. Data were obtained from GSE113764.[](G) Representative blots and quantification of SOX8 protein in corresponding human adipose tissues (n = 6).[](H) Correlation analysis between SOX8 mRNA levels and adipogenic differentiation of iBAT SVFs (n = 3).[]Data are presented as mean ± SEM. Statistical analysis was performed using Pearson correlation analysis (C and H), and unpaired two-tailed Student’s t-test (A, B and D-G). *p < 0.05, **p < 0.01.

We next examined SOX8 expression across different adipose depots. *Sox8* mRNA was not specifically enriched in mouse iBAT, as epididymal white adipose tissue (eWAT) also exhibited relatively high expression levels (Fig. S1A). Interestingly, SOX8 protein level was markedly enriched in iBAT (Fig. 1D), suggesting that SOX8 protein may serve as a specific marker of classic brown adipose tissue, potentially due to post-transcriptional regulation that limits its expression in white adipocytes. As adaptive thermogenesis activates under cold exposure, we next detected SOX8 expression after acute cold exposure. Although *Sox8* mRNA levels in iBAT did not change significantly (Fig.s S1B), SOX8 protein levels were markedly increased in iBAT after cold exposure in vivo (Fig. 1E). indicating that SOX8 is actively involved in adaptive thermogenesis in brown adipocytes. To assess whether this expression pattern is conserved in humans, we analyzed data from public human RNA-seq datasets and clinical samples. *SOX8* mRNA and protein were highly expressed in human brown adipose tissue (hBAT) (Fig.s 1F and 1G), supporting its role as a marker of human BAT. In addition, *Sox8* mRNA expression showed a significant increasing during brown adipogenic differentiation of iBAT-derived SVF cells (iBAT-SVFs) (Fig. 1H), showing SOX8 might affect adipogenesis.

These findings demonstrate that SOX8 is a conserved and specific marker of classic brown adipose tissue and associated with obesity. Mechanistically, SOX8 may promote thermogenesis through activation of the PGC-1α–UCP-1 axis, thereby representing a potential therapeutic target for obesity.

### SOX8 AKO mice lead to thermogenic dysfunction and obesity after HFD

To investigate the role of SOX8 in adipose tissue, we generated adipocyte-specific SOX8 knockout mice using Adipoq-Cre (AKO). *Sox8* mRNA levels were significantly decreased in the adipose tissues of AKO mice, confirming efficient gene deletion and validating the model (Fig.s S2A and S2B). Under normal diet (ND) conditions, no significant difference in total body weight was observed between AKO mice and control littermates (Fig.s S2C and S2D). However, iBAT and liver weights were increased in AKO mice (Fig.s S2E–S2H), suggesting early disturbances in lipid metabolism. Consistently, serum biochemical analyses revealed elevated alanine aminotransferase (ALT) and total cholesterol (TC) levels, accompanied by reduced triglyceride (TG) and free fatty acid (FFA) levels in AKO mice (Fig.s S2I–S2L), indicating liver damage, impaired lipid mobilization and metabolic imbalance under basal conditions. Impairments in glucose tolerance, insulin sensitivity, and cold tolerance were observed in ND-fed AKO mice (Fig.s S2M–S2O). Based on these observations, we subjected mice to a HFD feeding to further challenge metabolic homeostasis.

After only four weeks of HFD feeding, AKO mice displayed significantly greater body weight gain compared with control mice (Fig.s 2A and 2B). This increase was accompanied by a marked expansion of all adipose depots as well as a significant increase in liver weight, as reflected by both organ weight quantification and representative organ images (Fig.s 2C and 2D), indicating exacerbated lipid accumulation at both systemic and tissue levels. Interestingly, circulating FFA and TG levels were reduced in AKO mice (Fig.s 2E and 2F), suggesting impaired lipid mobilization and utilization, with lipids preferentially stored in adipose tissues. In contrast, low-density lipoprotein cholesterol (LDL-C) levels were elevated (Fig. 2G), indicating dysregulated lipid transport and an increased risk of cardiovascular complications. Histological analyses further supported these metabolic abnormalities. Adipocyte hypertrophy was evident across multiple adipose depots, including epididymal and inguinal white adipose tissue (Fig.s 2H and 2I). In parallel, hepatocyte enlargement was observed, together with enhanced hepatic steatosis (Fig.s 2H and 2J), suggesting that loss of SOX8 in adipose tissue exerts systemic metabolic consequences.

**Fig. 2.**
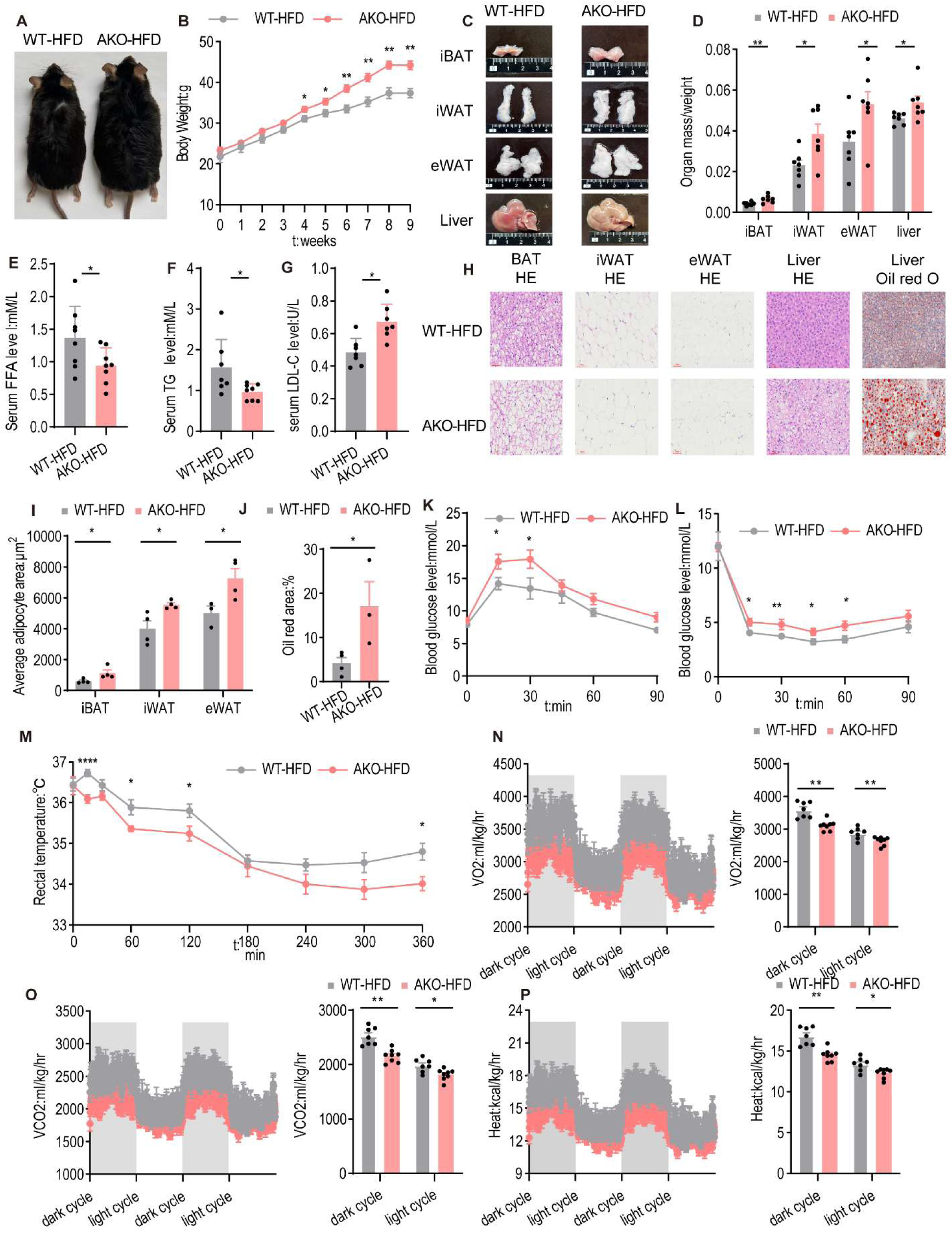
SOX8 AKO mice lead to thermogenic dysfunction and obesity after HFD.[](A) Representative images of HFD-fed mice.[](B) Body weight gain of mice fed a high-fat diet for 9 weeks (n = 7).[](C) Representative images of major organs from HFD-fed mice.[](D) Organ weights normalized to body weight in HFD-fed mice (n = 7).[](E-G) Serum triglyceride (TG), free fatty acid (FFA), and low-density lipoprotein cholesterol (LDL-C) levels in HFD-fed mice (n = 7-8).[](H) HE and Oil Red O staining of different tissues from HFD-fed mice.[](I) Quantification of adipocyte area in HFD-fed mice (n = 3-4).[](J) Quantification of lipid droplet area in the liver of HFD-fed mice (n = 4).[](K) Glucose tolerance test (GTT) in HFD-fed mice. Tail vein blood glucose levels were measured after 8 h fasting and intraperitoneal injection of glucose (2 mg/g body weight) (n = 7).[](L) Insulin tolerance test (ITT) in HFD-fed mice. Tail vein blood glucose levels were measured after 6 h fasting and intraperitoneal injection of recombinant human insulin (0.125 mg/g body weight) (n = 7).[](M) Cold tolerance test in HFD-fed mice. Rectal temperature was measured during cold exposure at 4°C after 6 h fasting (n = 7).[](N-P) Representative curves and quantification of oxygen consumption (VO₂), carbon dioxide production (VCO₂), and heat production in HFD-fed mice (n = 7).[]Data are presented as mean ± SEM. Statistical analysis was performed using an unpaired two-tailed Student’s t-test (B, D-G, I-P). *p < 0.05, **p < 0.01.

Similar to ND-fed mice, HFD-fed AKO mice exhibited significant impairments in glucose homeostasis and energy balance. Glucose tolerance tests (GTT) and insulin tolerance tests (ITT) revealed impaired glucose clearance and reduced insulin sensitivity (Fig.s 2K and 2L), indicating the development of insulin resistance and diabetic-like phenotypes. Furthermore, AKO mice showed a reduced ability to maintain body temperature during acute cold exposure (Fig. 2M), consistent with defective adaptive thermogenesis. To further characterize whole-body energy metabolism, we performed metabolic cage analyses at room temperature. Food and water intake were comparable between AKO and control mice (Fig.s S2P–S2Q), indicating that SOX8 deficiency does not affect feeding behavior or appetite. However, oxygen consumption (VO₂) and carbon dioxide production (VCO₂) were significantly decreased in AKO mice (Fig.s 2N–2O), suggesting reduced mitochondrial respiration and overall energy expenditure. Consistently, heat production was markedly diminished (Fig. 2P), further supporting impaired thermogenic capacity.

Collectively, these findings demonstrate that SOX8 is essential for maintaining adipose tissue metabolic homeostasis and adaptive thermogenesis. Loss of SOX8 leads to enhanced lipid storage, reduced energy expenditure, and systemic metabolic dysfunction, ultimately promoting obesity and insulin resistance. These results highlight SOX8 as a critical regulator of energy balance and a potential therapeutic target for obesity and related metabolic diseases.

### SOX8 regulates thePGC-1α-UCP-1/OXPHOS axis in mouse iBAT

To elucidate the molecular mechanism by which SOX8 regulates thermogenesis and confers resistance to obesity, we performed comprehensive RNA-seq and RT–qPCR analyses in mouse iBAT. Our analyses identified a total of 522 genes that were significantly altered in the iBAT of AKO mice (Fig. 3A). Subsequent pathway enrichment analysis revealed prominent changes in multiple metabolic pathways, with particularly pronounced alterations in those associated with mitochondrial function and overall energy metabolism (Fig. 3B). Notably, genes implicated in thermogenesis and oxidative phosphorylation (OXPHOS) were markedly downregulated, as demonstrated by RNA-seq quantification and RT-qPCR validation (Fig.s 3C–3F), collectively indicating a profound impairment in mitochondrial activity and a reduced capacity for heat production.

**Fig. 3.**
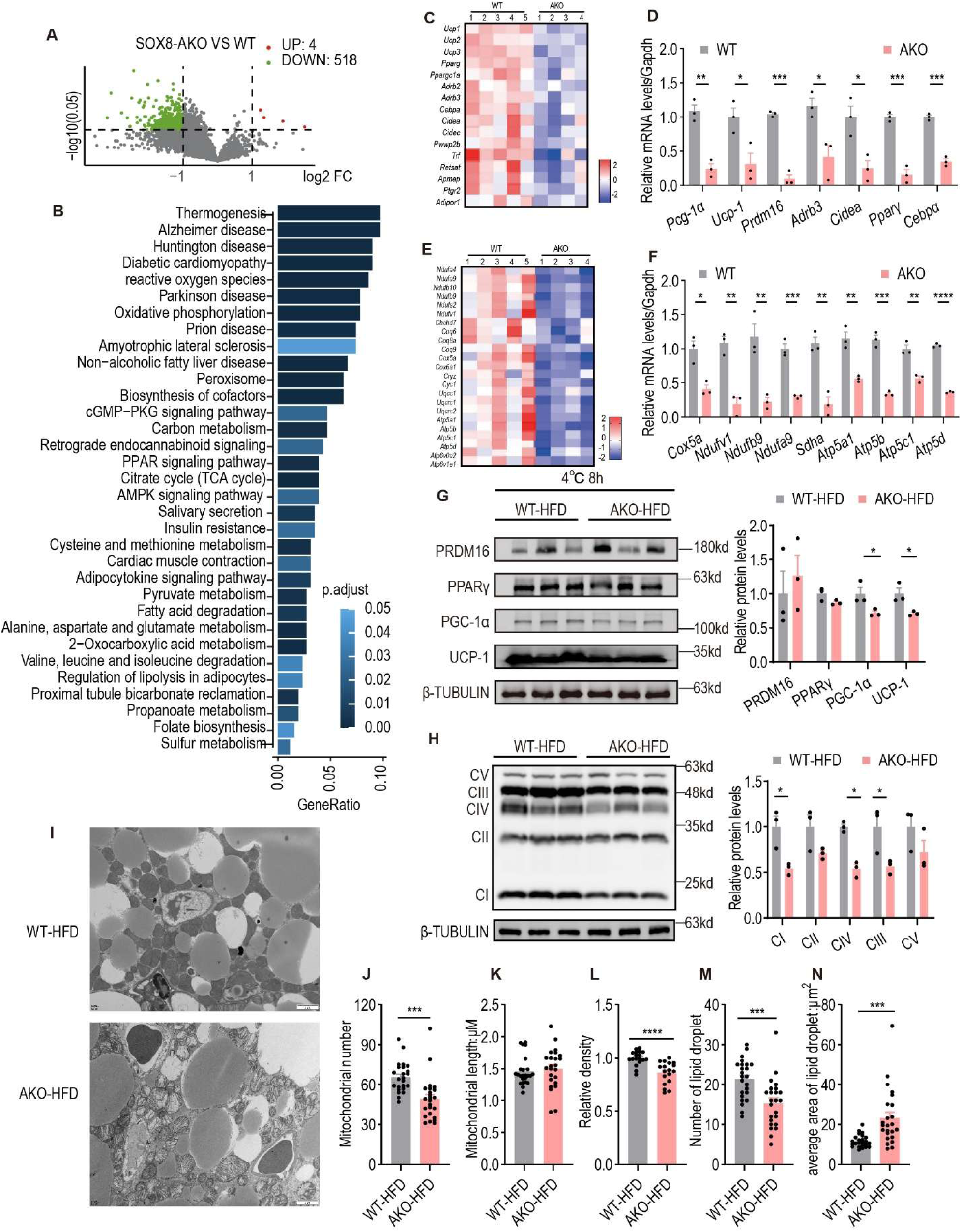
Loss of SOX8 leads to downregulation of thermogenesis and mitochondrial dysfunction in iBAT.[](A) Volcano plot showing significantly altered genes identified by RNA-seq in iBAT from AKO mice compared with controls (n = 4–5).[](B) Bar plot showing KEGG pathway enrichment analysis of differentially expressed genes from RNA-seq.[](C and D) Quantification of significantly altered thermogenic and OXPHOS gene expression levels identified by RNA-seq in AKO iBAT (n = 4–5).[](E and F) The mRNA expression of representative thermogenic genes and oxidative phosphorylation (OXPHOS)-related genes by RT-qPCR in iBAT (n = 3).[](G and H) Representative immunoblotting images and quantification of thermogenic and OXPHOS protein in iBAT from HFD-fed mice. After 6 h fasting, mice were exposed to 4°C for 6 h prior to tissue collection (n = 3).[](I) Representative transmission electron microscopy (TEM) images of interscapular brown adipose tissue (iBAT).[](J-N) Quantification of TEM images including mitochondrial number, mitochondrial length, mitochondrial density, lipid droplet number, and average lipid droplet area. (n = 18-24 from 3 mouse iBAT)[]Data are presented as mean ± SEM. Statistical analysis was performed using an unpaired two-tailed Student’s t-test (D, F-H and J-N). *p < 0.05, **p < 0.01, *** p < 0.001, ****p < 0.0001.

Further examination of adipogenic and thermogenic regulators revealed a striking divergence between transcriptional and protein-level regulation. While the mRNA levels of several adipogenic factors were significantly altered, the protein abundance of key adipogenesis regulators, including PRDM16 and PPARγ, remained largely unchanged (Fig. 3G), suggesting that SOX8 is largely dispensable for adipocyte differentiation per se. In contrast, thermogenesis-related proteins, including PGC-1α, UCP-1, and multiple components of the OXPHOS complexes, were substantially reduced, particularly under acute cold exposure conditions (Fig.s 3G and 3H). Given that PGC-1α serves as a master regulator of mitochondrial biogenesis and thermogenic gene expression, these results collectively suggest that SOX8 functions upstream of the PGC-1α–UCP-1/OXPHOS axis to modulate mitochondrial activity and adaptive thermogenesis.

Consistent with the observed suppression of OXPHOS signaling, transcriptomic analyses further demonstrated that SOX8 deficiency led to significant downregulation of genes involved in mitochondrial structural components as well as lipid metabolism (Fig.s S3A–S3C). These molecular alterations were corroborated at the ultrastructural level. Transmission electron microscopy (TEM) revealed a marked reduction in mitochondrial number, accompanied by disrupted cristae architecture, mitochondrial swelling, and enlarged lipid droplets in the iBAT of AKO mice (Fig.s 3I–3N). Such structural abnormalities are indicative of severe mitochondrial dysfunction and are frequently associated with reduced oxidative capacity and heightened metabolic stress within adipose tissue.

These findings establish SOX8 as a critical regulator for sustaining thermogenesis and promoting lipid utilization by modulating the PGC-1α–UCP-1/OXPHOS axis and preserving mitochondrial integrity. Importantly, our results highlight the central role of SOX8 in maintaining systemic energy balance and underscore its potential as a promising therapeutic target for combating obesity.

### SOX8 controls PGC-1α-dependent thermogenesis in brown adipocytes

SOX8 deficiency in adipose tissue led to impaired thermogenesis in vivo. To further investigate the underlying mechanisms, we next sought to determine whether SOX8 regulates adipocyte phenotypic remodeling, particularly the browning-to-whitening transition of brown adipocytes. Given the lack of specific markers for brown preadipocytes, we isolated the stromal vascular fraction from interscapular brown adipose tissue (iBAT-SVFs) of wild-type (WT) and SOX8 knockout (SOX8-KO) mice and immortalized these cells via stable expression of SV40 large T antigen. The resulting immortalized iBAT-SVFs were subsequently induced to differentiate into brown adipocytes (iBAs) for in vitro functional analyses (Fig.s S3D and S3E). Following adipogenic induction, SOX8-KO iBAs did not exhibit significant defects in overall adipocyte differentiation, consistent with our in vivo observations as these cells displayed increased lipid droplet accumulation (Fig. S4F), suggesting enhanced lipid storage and a partial shift toward a whitening phenotype. This observation indicates that while SOX8 is largely dispensable for the differentiation process itself, it plays a key role in maintaining brown adipocyte identity and limiting excessive lipid accumulation.

Functional analyses further revealed that the expression levels of key thermogenic and mitochondrial regulators, including PGC-1α and multiple components of the OXPHOS complexes, were markedly decreased in SOX8-KO iBAs following β3-adrenergic stimulation using CL316,243 (Fig.s 4A and 4B). Consistently, SOX8-deficient adipocytes exhibited a significantly reduced oxygen consumption rate (OCR) (Fig. 4C), reflecting compromised mitochondrial respiration and energy expenditure. These findings collectively suggest that SOX8 is essential for the proper activation of the thermogenic program and for sustaining mitochondrial function in brown adipocytes.

**Fig. 4.**
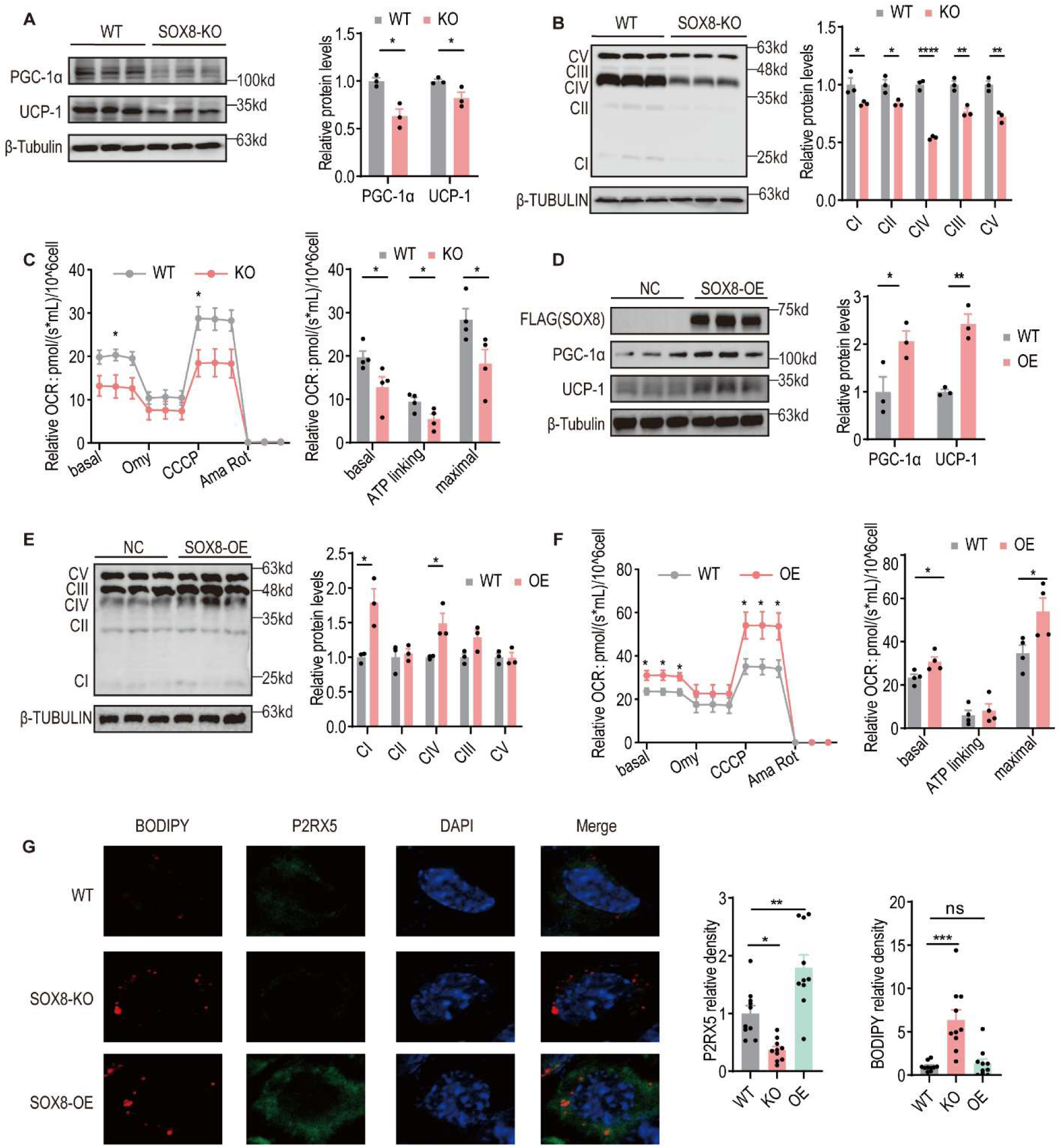
SOX8 controls PGC-1α-dependent thermogenesis in brown adipocytes.[](A and B) Representative western blots and quantification of thermogenic and OXPHOS protein levels of SVF-SOX8-KO-induced brown adipocytes (n = 3).[](C) Oxygen Consumption Curves and Quantitative Analysis of SVF-SOX8-KO-Induced Brown Adipocytes (n = 4).[](D and E) Representative western blots and quantification of thermogenic and OXPHOS protein levels of SVF-SOX8-OE-induced brown adipocytes (n = 3).[](F) Oxygen Consumption Curves and Quantitative Analysis of SVF-SOX8-OE-Induced Brown Adipocytes (n = 4).[](G) Representative immunofluorescence images and quantification of BODIPY staining and P2RX5 protein levels of SVF-induced brown adipocytes (n = 10)[]Data are presented as mean ± SEM. Statistical analysis was performed using an unpaired two-tailed Student’s t-test (A - F) and one-way analysis of variance (ANOVA) with Dunnett’s post multiple comparison test (G). nsp > 0.05, *p < 0.05, **p < 0.01, ***p < 0.001, ****p < 0.0001.

To determine whether SOX8 is sufficient to drive the thermogenic phenotype, we generated iBAs stably overexpressing SOX8 (SOX8-OE iBAs). Upon adipogenic induction, these cells displayed robust upregulation of thermogenic markers, including PGC-1α, UCP-1, and OXPHOS components (Fig.s 4D and 4E), supporting the reinforcement of brown adipocyte identity. Moreover, SOX8 overexpression significantly increased OCR (Fig. 4F), indicating enhanced mitochondrial respiration and energy expenditure. These results demonstrate that SOX8 is not only necessary but also sufficient to promote thermogenic competence in brown adipocytes.

Immunofluorescence staining further validated the effects of SOX8 manipulation on classic brown adipocyte identity, as reflected by changes in the brown adipocyte marker P2RX5(Ussar *et al*, 2014) in iBAs. Specifically, SOX8-KO led to a marked downregulation of P2RX5 accompanied by enlarged lipid droplets, collectively indicating a partial transition toward a white adipocyte-like phenotype (Fig. 4G). In contrast, SOX8-OE adipocytes exhibited elevated P2RX5 expression without concomitant lipid droplet accumulation (Fig. 4G). These observations suggest that SOX8 not only maintains brown adipocyte identity but also coordinates lipid storage and utilization, ensuring lipid homeostasis to support sustained thermogenic activity rather than merely suppressing lipid accumulation.

These findings provide compelling evidence that SOX8 is a critical regulator of brown adipocyte function. By maintaining mitochondrial activity and thermogenic capacity, SOX8 prevents pathological whitening of brown adipocytes, promotes energy expenditure, and coordinates the balance between lipid storage and utilization. SOX8 acts as a central node in the regulatory network controlling adipocyte phenotypic plasticity, thermogenic competency, and systemic energy homeostasis, highlighting its potential as a therapeutic target for combating obesity and metabolic disorders.

### SOX8 activates UCP-1 transcription and enhances PGC-1α stability through USP7

Having established that SOX8 regulates the expression of PGC-1α, UCP-1, and OXPHOS components, we next sought to elucidate the molecular mechanism by which SOX8 mediates thermogenesis in brown adipocytes. Considering that UCP-1 functions as the terminal effector of thermogenic activity, we first examined whether SOX8 directly regulates UCP-1 transcription. Consistent with changes observed at both the mRNA and protein levels, luciferase reporter assays revealed that SOX8 robustly enhanced the transcriptional activity of the UCP-1 promoter in both mouse and human brown adipocyte-derived stromal vascular fraction (BAT-SVF) cells (Fig.s S4A and S4B), suggesting a conserved regulatory mechanism across species. Chromatin immunoprecipitation (ChIP) assays further confirmed direct binding of SOX8 to the UCP-1 promoter region (Fig. S4C), establishing SOX8 as a bona fide transcriptional activator of UCP-1.

Given the central role of PGC-1α in mitochondrial biogenesis and thermogenic gene expression, we next investigated whether SOX8 regulates PGC-1α at the post-translational level. Cycloheximide (CHX) chase assays demonstrated that the half-life of PGC-1α was significantly reduced in SOX8-KO iBAs (Fig. 5A), indicating that SOX8 is required for maintaining PGC-1α protein stability. Consistently, treatment with the proteasome inhibitor MG132 prevented the downregulation of PGC-1α in SOX8-deficient cells, and K48-linked polyubiquitination of PGC-1α was markedly increased in SOX8-KO iBAs (Fig. 5B). These results collectively suggest that SOX8 stabilizes PGC-1α by suppressing K48-linked ubiquitin-mediated proteasomal degradation, thereby preserving the thermogenic capacity of brown adipocytes.

**Fig. 5.**
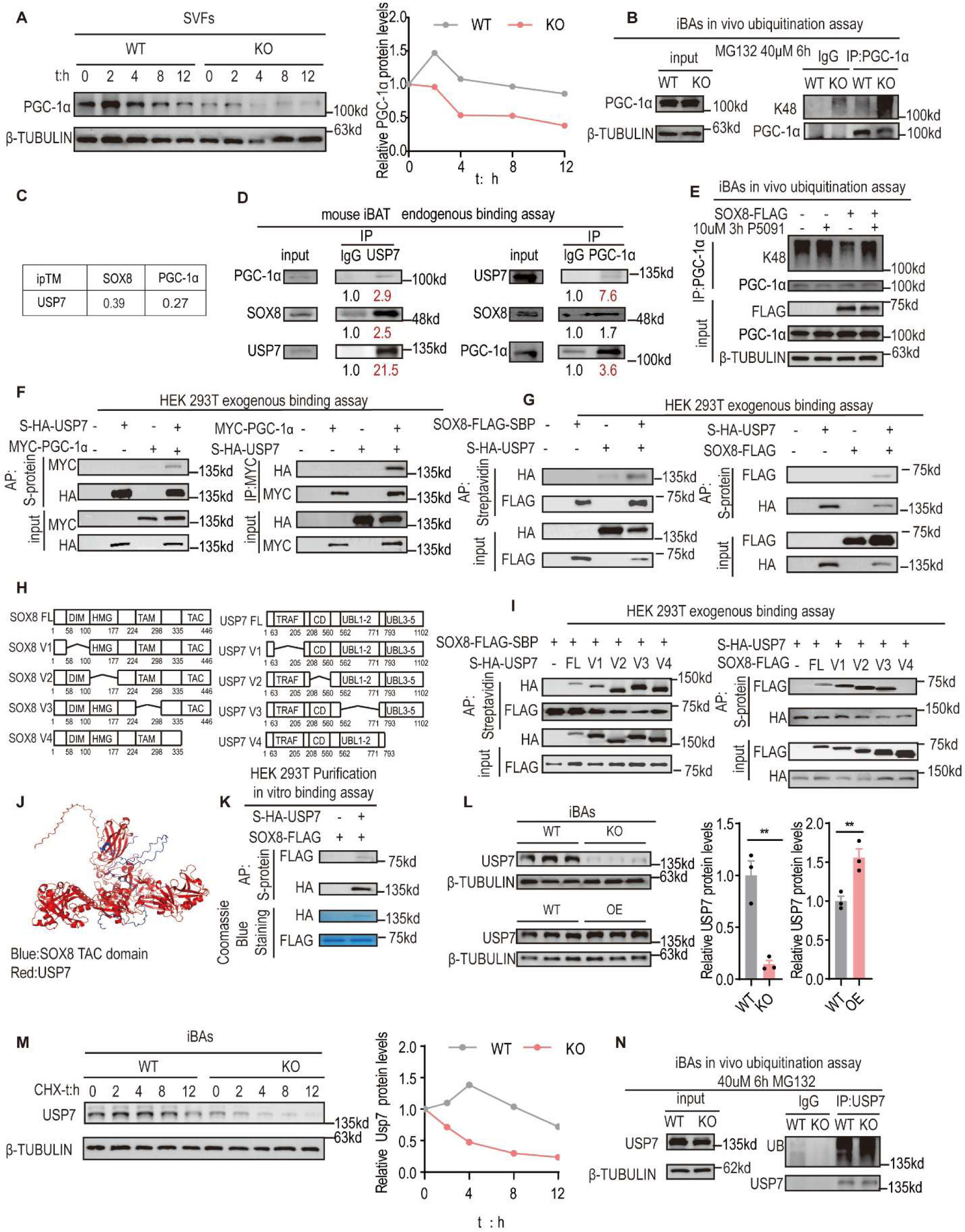
SOX8 activates UCP-1 transcription and PGC-1α stability through USP7.[](A) Representative blots and quantification of PGC-1α protein in SVFs subjected to a cycloheximide (CHX) chase assay.[](B) Representative blots of K48 ubiquitination levels of PGC-1α in SVF-derived brown adipocytes.[](C) AlphaFold3-predicted protein–protein interaction (PPI) scores for SOX8–USP7 and USP7–PGC-1α complexes.[](D) Representative blots and quantification of endogenous PPI among SOX8, USP7 and PGC-1α in mouse iBAT.[](E) Representative blots of K48-linked polyubiquitination levels of PGC-1α in SVF-induced brown adipocytes following SOX8 stable overexpression and treatment with the USP7 inhibitor P5091.[](F and G) Representative blots showing PPI between SOX8-USP7 and USP7-PGC-1α in HEK 293T cells after transfection with the indicated plasmids.[](H) Images for mutant construction of SOX8 and USP7.[](I) Representative blots of exogenous PPI among SOX8 and USP7 in HEK293T cells after transfecting indicated plasmids.[](J) AlphaFold3-predicted structural model of the SOX8 TAC domain and USP7 complex.[](K) Direct interaction between SOX8 and USP7 in vitro. Proteins were purified from HEK293T cells transfected with the indicated plasmids.[](L) Representative blots and quantification of USP7 protein in OE, WT and KO iBAs.[](M) Representative blots and quantification of USP7 protein in iBAs subjected to a CHX chase assay.[](N) Representative blots of USP7 ubiquitination levels in SVF-induced brown adipocytes.[]Data are presented as mean ± SEM. Statistical analysis was performed using an unpaired two-tailed Student’s t-test (N). *p < 0.05, **p < 0.01,*** p < 0.001,**** p < 0.0001.

Because SOX8 lacks intrinsic ubiquitin ligase or deubiquitinase activity, we hypothesized that it may stabilize PGC-1α through recruitment of a deubiquitinase (DUB). Structural predictions using AlphaFold3 identified ubiquitin-specific protease 7 (USP7) as a potential mediator capable of interacting with both SOX8 and PGC-1α (Fig. 5C). Co-immunoprecipitation (Co-IP) assays in mouse iBAT confirmed the existence of a SOX8–USP7–PGC-1α complex in vivo (Fig. 5D). To determine whether USP7 is the primary DUB mediating SOX8-dependent stabilization of PGC-1α, cells were treated with the USP7-specific inhibitor P5091. Inhibition of USP7 effectively abolished the ability of SOX8 overexpression to reduce K48-linked ubiquitination of PGC-1α, confirming that USP7 is a critical mediator linking SOX8 to PGC-1α stability (Fig. 5E).

To further dissect the interactions, co-transfection experiments in HEK293T cells revealed that SOX8 physically associates with USP7 (Fig. 5F), and USP7 in turn binds to PGC-1α (Fig. 5G). Interestingly, direct interaction between SOX8 and PGC-1α was not observed in HEK293T cells (Fig.s S4D and S4E), collectively indicating that USP7 functions as a molecular bridge connecting SOX8 to PGC-1α. Purified protein assays confirmed that SOX8 enhances USP7-mediated removal of K48-linked polyubiquitin chains from PGC-1α in vitro (Fig. S4F), supporting a model in which SOX8 recruits USP7 to its substrates. Next, we mapped the structural determinants of the SOX8–USP7 interaction. Truncation analyses revealed that the TAC (transactivation C-terminal) domain of SOX8 is necessary for USP7 binding, whereas deletion of individual USP7 domains did not completely abolish the interaction (Fig.s 5H and 5I), suggesting that SOX8 may engage multiple regions of USP7 or stabilize its conformation through a noncanonical binding mode. Consistent with this, AlphaFold3 structural modeling showed that the TAC domain of SOX8 wraps around the interior of USP7 in a spring-like manner, forming a distinct binding pocket (Fig. 5J). In vitro binding assays using purified proteins further validated the direct interaction between SOX8 and USP7 (Fig. 5K), indicating that SOX8 not only serves as a scaffold but also potentially enhances USP7 substrate accessibility or enzymatic activity. In addition to facilitating complex formation, SOX8 also regulates USP7 protein abundance and stability. SOX8 knockout markedly reduced USP7 levels, whereas SOX8 overexpression increased USP7 abundance (Fig. 5L). CHX chase assays demonstrated that SOX8 deficiency shortened the half-life of USP7 (Fig. 5M), accompanied by increased ubiquitination (Fig. 5N). Interestingly, MG132 treatment did not restore USP7 protein levels in SOX8-deficient cells and total ubiquitination of USP7 was markedly increased in SOX8-KO iBAs (Fig. 5N), suggesting that USP7 degradation is not primarily proteasome-dependent. In vitro deubiquitination assays further revealed that SOX8 reduces K63-linked polyubiquitination of USP7 (Fig. S4G), implying regulation via the autophagy–lysosome pathway.

Collectively, these results delineate a feed-forward regulatory axis in brown adipocytes, in which SOX8 simultaneously promotes UCP-1 transcription, recruits and activates USP7, and stabilizes PGC-1α protein. In parallel, SOX8 preserves USP7 abundance, ensuring sustained deubiquitinase activity. This dual regulatory role of SOX8 reinforces thermogenic gene expression, mitochondrial function, and energy expenditure, highlighting the SOX8–USP7–PGC-1α axis as a central pathway for adaptive thermogenesis and a potential therapeutic target for obesity intervention.

### AAV-mediated SOX8 overexpression exerts time-dependent effects on thermogenesis and metabolic homeostasis in HFD-fed mice

To investigate the therapeutic potential of SOX8 in vivo, we overexpressed SOX8 in adipose tissue using AAV-Rec2-mediated gene delivery(Huang *et al*, 2016) and fed on HFD for 6 weeks. Although body weight remained largely unchanged compared with control mice (Fig. 6A), iBAT weight was significantly reduced (Fig. 6B), and histological analyses revealed smaller adipocyte size and more compact tissue architecture (Fig. 6C), indicating enhanced lipid utilization and BAT remodeling. Consistently, fasting blood glucose levels were lower in SOX8-overexpressing mice, and insulin tolerance tests demonstrated improved insulin sensitivity (Fig.s 6D and 6E), suggesting that SOX8 promotes systemic glucose homeostasis. Rectal temperature measurements showed a significant increase in basal thermogenesis (Fig. 6F), which was further supported by metabolic cage analyses demonstrating elevated oxygen consumption (VO₂), carbon dioxide production (VCO₂), and heat production (Fig.s 6G–6I) without food or water intake difference (Fig.s S5A and S5B). These findings collectively indicate that SOX8 enhances whole-body energy expenditure.

**Fig. 6.**
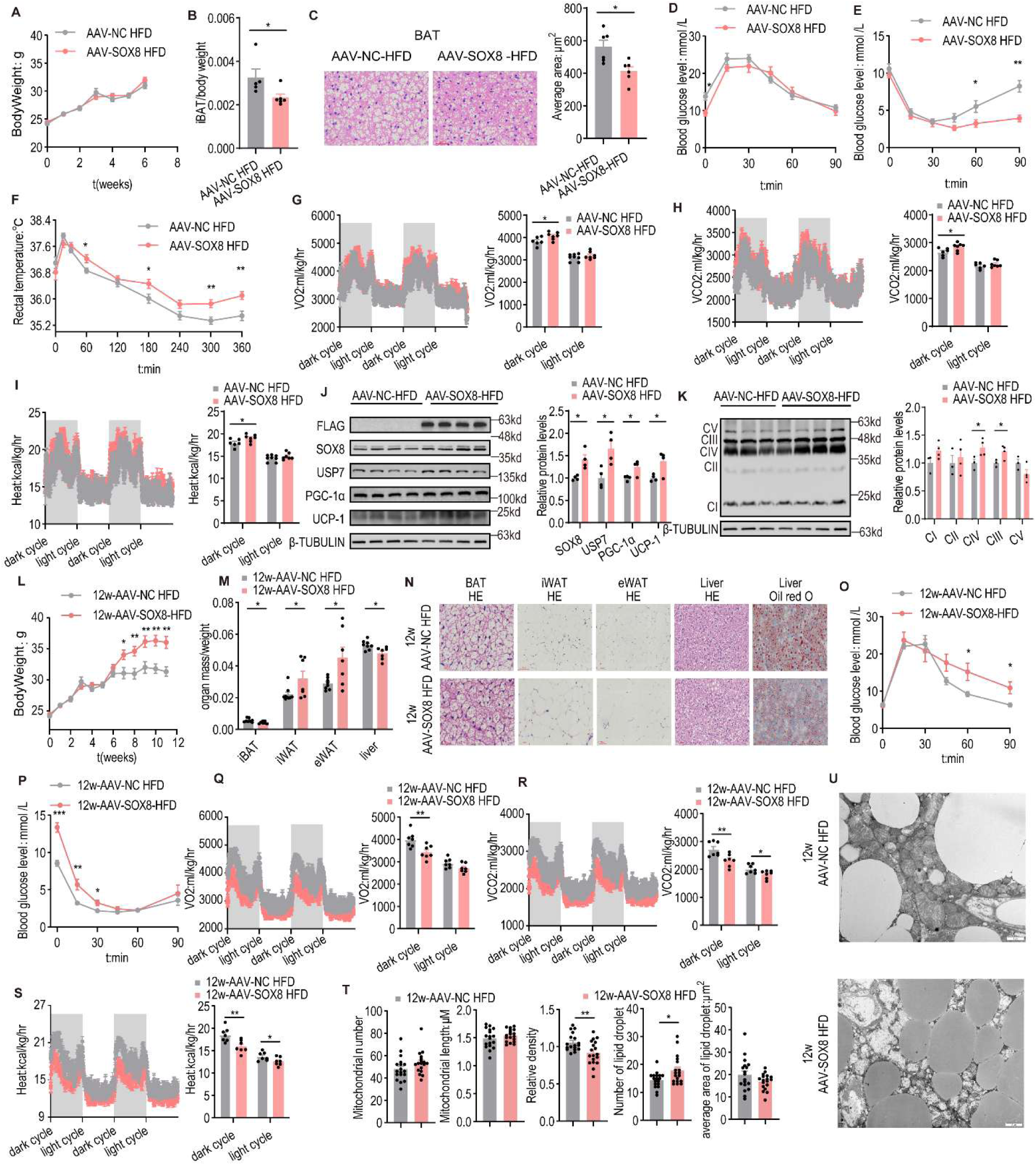
AAV-mediated SOX8 overexpression exerts time-dependent effects on thermogenesis and metabolic homeostasis in HFD-fed mice.[](A) Body weight gain in mice fed a high-fat diet for 6 weeks (n = 6).[](B) The iBAT weight normalized to body weight in 6-week HFD-fed mice (n = 6).[](C) Representative HE staining of iBAT and quantification of adipocyte area from AAV-treated 6-week HFD-fed mice (n = 6).[](D) Glucose tolerance test (GTT) in AAV-treated 6-week HFD-fed mice. Tail vein blood glucose levels were measured after 8 h fasting followed by intraperitoneal injection of glucose (2 mg/g body weight) (n = 6).[](E) Insulin tolerance test (ITT) in AAV-treated 6-week HFD-fed mice. Tail vein blood glucose levels were measured after 6 h fasting followed by intraperitoneal injection of recombinant human insulin (0.125 mg/g body weight) (n = 6).[](F) Cold tolerance test in AAV-treated 6-week HFD-fed mice. Rectal temperature was recorded during cold exposure at 4°C after 6 h fasting (n = 6).[](G–I) Representative curves and quantification of oxygen consumption (VO₂), carbon dioxide production (VCO₂), and heat production in AAV-treated 6-week HFD-fed mice (n = 6).[](J–K) Representative immunoblotting images and quantification of indicated proteins in iBAT from AAV-treated 6-week HFD-fed mice.[](L) Body weight gain in mice fed a high-fat diet for 12 weeks (n = 9).[](M) Organ weights normalized to body weight in 12-week HFD-fed mice (n = 7-9)[](N) HE and Oil Red O staining of different tissues from 12-week HFD-fed mice.[](O) Glucose tolerance test (GTT) in AAV-treated 12-week HFD-fed mice. Tail vein blood glucose levels were measured after 8 h fasting followed by intraperitoneal injection of glucose (2 mg/g body weight) (n = 7).[](P) Insulin tolerance test (ITT) in AAV-treated 12-week HFD-fed mice. Tail vein blood glucose levels were measured after 6 h fasting followed by intraperitoneal injection of recombinant human insulin (0.125 mg/g body weight) (n = 9).[](Q–S) Representative curves and quantification of oxygen consumption (VO₂), carbon dioxide production (VCO₂), and heat production in AAV-treated 12-week HFD-fed mice (n = 7).[](T) Quantification of TEM images including mitochondrial number, mitochondrial length, mitochondrial density, lipid droplet number, and average lipid droplet area from 12-week HFD-fed mice. (n = 18 from 3 mouse iBAT)[](U) Representative transmission electron microscopy (TEM) images of interscapular brown adipose tissue (iBAT) from 12-week HFD-fed mice.[]Data are presented as mean ± SEM. Statistical significance was determined using an unpaired two-tailed Student’s t-test (A-K). *P < 0.05, **P < 0.01, *** p < 0.001.

At the molecular level, AAV-mediated SOX8 overexpression led to markedly increased protein levels of FLAG-tagged SOX8, PGC-1α, UCP-1, and components of OXPHOS machinery in iBAT (Fig.s 6J and 6K). The coordinated upregulation of these thermogenic and mitochondrial proteins suggests that SOX8 activates the PGC-1α–UCP-1/OXPHOS axis and promotes mitochondrial biogenesis, thereby enhancing the thermogenic capacity of brown adipocytes.

Under ND conditions, adipose-specific overexpression of SOX8 did not significantly affect body weight, organ weights, adipocyte morphology, glucose tolerance, cold tolerance, energy expenditure, or thermogenic protein expression (Fig.s S5C–S5F and S5H–S5O). However, insulin sensitivity was modestly improved, as indicated by enhanced insulin tolerance (Fig. S5G). These results suggest that SOX8 exerts minimal effects on basal metabolic homeostasis but confers metabolic benefits under conditions of metabolic stress.

To assess the long-term effects of sustained SOX8 overexpression, we extended the HFD intervention period to 12 weeks. In contrast to the beneficial metabolic effects observed during short-term overexpression, prolonged SOX8 activation resulted in significant body weight gain compared with controls (Fig. 6L), suggesting the emergence of compensatory mechanisms that progressively attenuate its metabolic benefits. Organ weight analyses revealed significantly increased iWAT and eWAT mass, whereas iBAT and liver weights were reduced in SOX8-overexpressing mice (Fig. 6M). Histological examination showed a trend toward reduced adipocyte size in BAT, accompanied by adipocyte hypertrophy in iWAT and eWAT, although these changes did not reach statistical significance (Fig.s 6N and S5P). H&E and Oil Red O staining further demonstrated no overt hepatic injury and modestly reduced lipid accumulation in the livers of long-term SOX8-overexpressing mice (Fig.s 6N and S5Q), indicating distinct tissue-specific metabolic effects following prolonged SOX8 activation. Notably, whereas short-term SOX8 overexpression improved insulin sensitivity, prolonged overexpression markedly impaired both glucose tolerance and insulin sensitivity (Fig.s 6O and 6P), indicating the emergence of systemic metabolic dysfunction with extended intervention.

Further assessment of thermogenesis and energy metabolism revealed that cold tolerance was comparable between long-term SOX8-overexpressing mice and controls during acute cold exposure at 4°C (Fig. S5R), suggesting that chronic SOX8 overexpression does not significantly impair cold-adaptive thermogenesis.

However, despite comparable food and water intake (Fig.s S5S and S5T), metabolic cage analyses revealed significantly reduced oxygen consumption (VO₂), carbon dioxide production (VCO₂), and heat production in SOX8-overexpressing mice (Fig.s 6Q–6S), indicating a pronounced decline in whole-body energy expenditure. These findings suggest that chronic SOX8 activation induces compensatory suppression of thermogenic activity, ultimately leading to systemic metabolic maladaptation despite partial preservation of BAT morphology. Consistent with this functional deterioration, TEM analysis of iBAT revealed severe mitochondrial ultrastructural abnormalities in long-term SOX8-overexpressing mice. Although mitochondrial number remained unchanged, mitochondrial cristae were largely disrupted and accompanied by increased lipid droplet accumulation (Fig.s 6T and 6U), indicating profound mitochondrial dysfunction following chronic SOX8 activation.

Collectively, these findings demonstrate that the metabolic effects of SOX8 overexpression are highly time-dependent. Whereas short-term activation enhances BAT thermogenesis and improves systemic metabolic homeostasis, prolonged overexpression paradoxically induces compensatory metabolic deterioration. These results highlight the importance of temporally controlled SOX8-targeted interventions for the treatment of obesity and metabolic disorders.

## Discussion

Obesity is driven by chronic imbalance between energy intake and expenditure, and impaired brown adipose tissue (BAT) thermogenesis has emerged as a major contributor to metabolic dysfunction(Cannon & Nedergaard, 2004). Increasing energy expenditure through BAT activation is therefore considered a promising therapeutic strategy for obesity and related metabolic diseases (Elmaleh-Sachs *et al*, 2023; Hinney *et al*, 2022; Perdomo *et al*., 2023). In this study, we identified SOX8 as a previously unrecognized BAT-enriched thermogenic regulator that maintains mitochondrial integrity and systemic metabolic homeostasis through the USP7–PGC-1α axis.Previous studies have mainly focused on transcription factors regulating adipocyte differentiation and thermogenic gene expression, such as PPARγ, PRDM16, and PGC-1α (Lin *et al*, 2005a; Ohno *et al*, 2012). However, recent investigations have provided new insights into how to coordinately integrate thermogenic program activation and mitochondrial homeostasis at the upstream transcriptional level to achieve metabolic remodeling and obesity control. Specifically, the BAT-specific transcription factor EBF2 recruits PPARγ to the regulatory regions of BAT-specific genes(Rajakumari *et al*, 2013), thereby driving brown and beige adipocyte differentiation and activating thermogenic gene expression. Moreover, the nuclear receptor co-regulator NR2F6 in brown preadipocytes promotes PPARγ expression and facilitates brown adipocyte differentiation(Zhou *et al*, 2024). Additionally, knockout of MTCH2 enhances BAT thermogenesis and subcutaneous white fat browning, protecting against HFD-induced obesity, while deletion of the transcription factor JunB alters the brown adipocyte phenotype, increases cellular thermogenic capacity, and improves metabolic function(Zhang *et al*, 2024; Zhao *et al*, 2025).

SOX8 is recently involved in ear and M cell differentiation(Buzzi *et al*, 2022; Kimura *et al*, 2019). Previous studies have identified SOX8 expression in metabolically relevant tissues, including mesodermal regions destined for adipogenesis and in undifferentiated mouse embryonic fibroblasts (MEFs)(González Alvarado & Aprato, 2025; Guth *et al*., 2009). These observations suggest that SOX8 is a novel transcription factor with regulatory functions in adipocyte development. Early in vivo studies further demonstrate that global SOX8 knockout in mice resulted in mild reductions in bone mass and adipose tissue mass in adulthood, leading to decreased body weight(Guth *et al*., 2009; Stolt *et al*, 2004). Our previous work reveals that SOX8 regulated de novo lipogenesis, glucose metabolism, and mitochondrial structure and function(Yang *et al*, 2024b). The present study extended these findings by demonstrating that SOX8 was highly and specifically expressed in BAT, and acted as a novel thermogenic-responsive transcriptional regulator. Under cold exposure, SOX8 expression dynamics were highly synchronized with those of the classic thermogenic regulators PGC-1α and UCP-1. Specific ablation of SOX8 in adipose tissue induced a pronounced “whitening” phenotype in BAT, characterized by reduced multilocular small lipid droplets, increased unilocular large lipid droplets, and enlarged cell size, indicating a critical role for SOX8 in maintaining brown adipocyte identity and homeostasis.

Furthermore, adipose-specific SOX8 knockout (AKO) mice exhibited severe metabolic disturbances, including fatty liver, glucose intolerance, and insulin resistance, all of which were indicative of systemic energy homeostasis disruption. Notably, the metabolic phenotype of AKO mice was highly diet-dependent: body weight remained unchanged under a chow diet, but increased significantly following HFD feeding. These suggest that under normal dietary conditions, the mild impairment of BAT thermogenic function caused by SOX8 deficiency could be compensated for by alternative mechanisms to maintain energy balance. In contrast, HFD-induced metabolic stress led to energy overload that exceed compensatory capacity, and AKO mice fell to efficiently activate thermogenesis to dissipate excess energy, resulting in overt obesity. This phenomenon is consistent with previous findings in metabolic research: adipose-specific Gys1 knockout mice show no significant differences in body weight or body composition under a chow diet, but develop marked metabolic abnormalities under HFD feeding(Zhuo *et al*, 2024). Similarly, preadipocyte-specific NR2F6 knockout mice exhibit no significant difference in body weight on a chow diet but develop severe obesity, glucose intolerance, insulin resistance, and fatty liver after 12 weeks of HFD feeding(Zhou *et al*., 2024). Serum biochemical parameters also exhibited distinct patterns under the two dietary conditions. Reduced serum free fatty acid (FFA) levels in AKO mice were likely attributed to impaired BAT thermogenic function following SOX8 ablation, as thermogenesis required extensive fatty acid mobilization; thus, diminished lipolysis directly reduced FFA release from adipose tissue. Consistent with this, adipose-specific Sirt6 knockout mice also exhibit impaired lipolytic activity due to decreased ATGL expression. As the rate-limiting enzyme in lipolysis, reduced ATGL expression directly decreases intracellular triglyceride hydrolysis and FFA release(Kuang *et al*, 2017). Since the present study did not directly measure the expression of lipolytic enzymes such as ATGL and HSL, this interpretation awaits further experimental validation. When the balance between lipid storage and release in adipose tissue was disrupted, excess lipids might ectopically accumulate in the liver, perturb hepatic cholesterol metabolism, and thereby led to elevated serum total cholesterol (TC) levels and increased LDL-C under HFD conditions. These diet-dependent differences reflect the dynamic progression of adipose tissue dysfunction caused by SOX8 deficiency.

Mitochondria serve as the ultimate effectors of thermogenesis, and their structural and functional integrity is a key determinant of BAT thermogenic capacity. Transmission electron microscopy in the present study revealed that SOX8 deficiency induced mitochondrial structural abnormalities, including swelling and cristae loss, which were further exacerbated by HFD feeding. Under a chow diet, mitochondria in AKO mice were shorter, whereas under HFD, they became elongated, suggesting differential dynamics of mitochondrial fusion and fission under varying intensities of metabolic stress, with the underlying mechanisms requiring further investigation. Recent studies have also highlighted the critical role of intact mitochondrial cristae architecture in thermogenic function: for example, ELOVL3 deficiency impairs mitochondrial cristae remodeling and UCP-1 expression in BAT(Westerberg *et al*, 2006), and adipose-specific Tmem135 knockout blocks mitochondrial fission, impairs thermogenesis, and exacerbates diet-induced obesity(Hu *et al*, 2023), whose findings that are consistent with our observations. Using iBAT-SVF cells induced to differentiate into brown adipocytes in vitro, we further demonstrated that SOX8-deficient cells exhibited significantly reduced basal respiration, ATP-coupled respiration, and maximal respiratory capacity, confirming a cell-autonomous, positive regulatory role of SOX8 in mitochondrial function. Collectively, mitochondrial dysfunction leads to incomplete fatty acid β-oxidation, electron leakage, and excessive reactive oxygen species (ROS) production; ROS accumulation further impairs insulin signaling, ultimately driving adipocyte dysfunction(Furukawa *et al*, 2004). Our study thus establishes that SOX8 played a critical cell-autonomous role in maintaining mitochondrial morphology and functional stability in BAT.

Mechanistically, SOX8 functioned as a multifaceted regulator. It transcriptionally enhanced UCP-1 expression, reinforcing thermogenic effector programs, while simultaneously stabilizing PGC-1α protein by recruiting USP7, which removed K48-linked ubiquitin chains and prevent proteasomal degradation. Notably, SOX8 maintained USP7 protein abundance and appeared to modulate its enzymatic activity, likely through a specific interaction interface within the high-mobility group (HMG) domain or the TAC domain of SOX8.

These structural elements might facilitate conformational changes in USP7 that enhanced its deubiquitinase activity toward PGC-1 coactivators and other substrates, forming a feed-forward regulatory loop that sustains adaptive thermogenesis. USP7 itself also influences adipocyte differentiation, affecting lineage commitment, PPARγ stability, lipid droplet formation, and thermogenic programming. Besides, in brown adipocytes induced by iBAT-SVF in vitro differentiation, SOX8 knockout significantly reduced the ratio of mature brown adipocytes, whereas SOX8 overexpression increased the number of brown adipocytes. We suspect that SOX8 differentially modulated USP7 levels in preadipocytes versus mature adipocytes, where SOX8 regulated the substrate specificity of USP7 depending on the differentiation stage, which further highlights its dual role in coordinating adipocyte development and mature brown adipocyte function. Furthermore, USP7 interacts with RNF2(Yan *et al*, 2025), an E3 ligase mediating PGC-1α ubiquitination(Sen *et al*., 2011); USP7 also interplays with p62(Chen *et al*, 2025), a selective autophagy receptor. Beyond the canonical SOX8–USP7–PGC-1α axis, we suspected that there might be multi-layered regulatory networks, where SOX8 integrated USP7–RNF2–PGC-1α or USP7–p62 regulatory pathways, ensuring sustained thermogenic capacity, mitochondrial integrity, and systemic metabolic homeostasis.

From a therapeutic perspective, SOX8 represents a promising target for the treatment of obesity and metabolic disorders. By using AAV-Rec2-mediated gene delivery to maintain elevated SOX8 expression in brown adipocytes, our study uncovered a previously unrecognized time-dependent effect of SOX8 activation on systemic metabolism. Short-term SOX8 overexpression (6 weeks) markedly improved metabolic phenotypes in HFD-fed mice, as evidenced by reduced brown adipocyte size, increased oxygen consumption and core body temperature, and enhanced insulin sensitivity, supporting the feasibility of targeting SOX8 to enhance BAT thermogenesis and combat obesity.

However, when the intervention period was extended to 12 weeks, the metabolic phenotype was completely reversed. Prolonged SOX8 overexpression resulted in compensatory body weight gain, exacerbated glucose intolerance and insulin resistance, and a pronounced decline in whole-body energy expenditure compared with control mice. In the field of metabolic research, this phenomenon is commonly referred to as metabolic maladaptation. One possible explanation for this paradoxical effect is that chronic SOX8 activation continuously stabilizes PGC-1α protein, thereby maintaining mitochondria and UCP1-dependent thermogenesis in a persistently hyperactivated state. Sustained mitochondrial overactivation may lead to excessive reactive oxygen species (ROS) accumulation, progressive oxidative damage, and activation of negative feedback mechanisms that suppress energy expenditure in order to preserve systemic energy homeostasis. Consistent with this hypothesis, long-term SOX8-overexpressing mice exhibited severe mitochondrial ultrastructural disruption and reduced thermogenic activity despite partial preservation of BAT morphology.

Importantly, similar time-dependent effects have also been reported in studies involving chronic activation of the β-adrenergic signaling pathway. Although prolonged administration of the β3-adrenergic receptor (β3-AR) agonist CL316243 initially promotes lipid utilization and thermogenesis, sustained stimulation can induce attenuation of β-adrenergic responsiveness and adaptive tissue remodeling, thereby progressively diminishing its anti-obesity efficacy (Ferrand *et al*, 2006). In addition, transgenic overexpression of the mitochondrial outer membrane protein mitoNEET in adipose tissue was previously shown to promote thermogenic activation during the early stages of HFD feeding; however, after prolonged feeding, PGC-1α and UCP1 expression became markedly suppressed, followed by rapid obesity development(Kusminski *et al*, 2014). Together, these findings suggest that chronic overstimulation of thermogenic pathways may ultimately trigger adverse adaptive reprogramming of metabolic systems. Therefore, our findings not only reveal a time-dependent role of SOX8 in regulating BAT thermogenesis and systemic energy homeostasis, but also provide important implications for the development of BAT-targeted therapies. Interventions aimed at activating SOX8 or other thermogenic regulators may require precise temporal and dosage control to maximize metabolic benefits while avoiding chronic mitochondrial stress and maladaptive compensatory responses.

In conclusion, the SOX8-USP7-PGC-1α axis provided a mechanistic framework for understanding how adipocyte-intrinsic networks control energy balance, substrate utilization, and cell survival, which underscores its potential to combat obesity and metabolic dysfunction.

## CRediT authorship contribution statement

Conceptualization, **H Q**and **ZY W**; methodology, **YQ G** and **ZH K**; Investigation, **YQ G**, **ZH K**, **GY L**, **XJ Y**, **Q Z** and **YW L, X H, X Y**; writing - original draft, **YQ G** and **ZH K**; writing - review and editing, **ZY W**; funding acquisition, **ZL Y**, **H Q** and **ZY W**; resources, **YC C**, **ZL Y** and **ZY W**; supervision, **H Q** and **ZY W**.

## Acknowledgements

We are grateful to patients for donating their adipose tissues.

## Appendix A. Supplementary data

Supplementary data to this article can be found in PDF entitled “Appendix A. Supplementary data”.

## Data availability

All data generated or analyzed during this study are available from the corresponding authors upon reasonable request. The RNA-seq data have been deposited in the National Genomics Data Center (NGDC) under accession number OMIX016363 and are publicly available at: https://ngdc.cncb.ac.cn/omix/release/OMIX016363

